# Pharmacodynamic profiles inform systematization, efficacy and whole-brain drug distribution profiles in 18 antidepressants

**DOI:** 10.64898/2026.04.26.720902

**Authors:** Benjamin Hänisch, Sofie L. Valk

**Author notes:** Correspondence: Benjamin Hänisch: Department of Psychiatry and Psychotherapy, University Hospital Tübingen, Tübingen, Germany., Sofie L. Valk: Max Planck Institute for Human Cognitive and Brain Sciences, Leipzig, Germany.

## Abstract

Although empirical evidence shows that antidepressants are effective and superior to placebo in treating depressive disorders, their mechanisms of action are still unclear. In this study, we examine the multidimensional molecular affinities of 18 commonly used antidepressants. Clustering analyses consistently generate three distinct clusters, providing a broader taxonomy that groups SSRIs and SNRIs together. Correlational analyses cautiously indicate a relationship between the affinity to metabotropic serotonin receptors and antidepressant efficacy. Next, we generate anatomical distribution profiles of drug action strengths by combining positron emission tomography-derived maps of cerebral neurotransmitter receptor and transporter densities from an open-access repository of healthy participants with the drugs’ affinity profiles. We then relate these profiles to functional and structural neuroanatomical measures in health and disease. Our results reveal distinct, mechanistically interpretable differences between antidepressants with high 5-HTT affinity and atypical antidepressants. These differences could inform personalized drug selection and development.

## Introduction

Depressive disorders are major contributors to the global burden of disease (1), and antidepressant medication is an important constituent of guideline-based care for major depressive disorder. Most antidepressants inhibit transporters of monoamine neurotransmitters, increasing the concentration of neurotransmitters in the synaptic cleft, while others directly bind metabotropic neurotransmitter receptors, mostly from the serotonin and noradrenaline systems. Even though the clinical efficacy of antidepressants is well-established, and their specific inhibitory properties are known, how pharmacological neuromodulation alleviates depressive symptoms is still unclear. Nevertheless, there are several theories that try to explain the phenomenon. The prominent monoamine hypothesis states that depressive symptoms are caused by a depletion of monoaminergic neurotransmitters, which gets replenished through antidepressant-mediated transporter inhibition (2). This hypothesis could not be verified despite considerable scientific efforts (3). More recent studies support the neuroplasticity hypothesis, which states that induction of neuroplastic effects through antidepressant medication is vital to mediate an improvement of depressive symptoms (4,5). On a neuropsychological level, proposals regarding their alleviation of a negative bias inherent in processing emotionally salient stimuli have recently gained traction (6). Despite the existence of different theories with varying levels of empirical support, however, uncovering a clear mechanism of action that links neurotransmitter system perturbation to depressive symptom alleviation has thus far not been satisfactorily achieved.

The lack of understanding regarding the relationship between pharmacodynamic profiles and the varying clinical efficacy of different antidepressant drugs (7) impairs both developments of novel antidepressants and more patient-specific choices of antidepressant treatment. This - next to the unclear pathophysiology underlying a depressive syndrome (8) - presents a major roadblock to reaching the goal of precision medicine in psychiatry (9). Furthermore, a neurobiologically plausible explanation why antidepressants are also effective in treating other psychiatric conditions, such as anxiety disorder and obsessive-compulsive disorder, remains similarly elusive (10,11). Evidence from neuroplasticity studies suggests that antidepressant treatment response is mediated through different pathways than disease onset (12), opening up the possibility for a common down-stream effect of psychopharmacological treatment with antidepressant drugs across multiple psychiatric conditions. Summarized, despite considerable research efforts since their serendipitous discovery several decades ago, central questions regarding the mechanisms of action through which antidepressants influence psychopathological states remain unanswered.

To gain further insights into the neurobiological mechanism of action of antidepressant drugs, this study follows a two-step approach. We first focus on the relationship between pharmacodynamic properties and antidepressant efficacy. Both our choice of antidepressant substances and our measure of clinical efficacy are based on the results of a recent network meta-analysis by Cipriani and colleagues, which showed varying degrees of antidepressant efficacies across 21 substances that were all superior to placebo controls (7). Interestingly, the traditional antidepressant systematization into different reuptake inhibitors, tricyclic and atypical antidepressants does not clearly inform on the variations in clinical efficacy. This warrants a more detailed study of antidepressants’ pharmacodynamic profiles, which we approach through generating substance-specific multidimensional affinity profiles that allow for data-driven systematization techniques. To assess molecular affinities, we use inhibition constant (Ki) values, a pharmacodynamic measure that quantifies inhibitory potency in substrate-ligand interactions. We derive these Ki values from the Psychoactive Drug Screening Program Ki Database, and open-access collection of experimentally established Ki values (KiDB) (13). In a second step, we use these affinity profiles to develop a neuroanatomical perspective on antidepressant action. As neuroanatomical studies have yielded considerable insights into the structure-function relationships in the human brain, we seek to leverage the strength and detail of neuroanatomical knowledge to establish a new perspective on antidepressant drugs. We combine antidepressants’ affinity profiles with Positron Emission Tomography (PET)-derived neurotransmitter transporter and receptor density maps from an open-access repository of healthy participants (14), to describe anatomical drug distribution profiles in the human cerebral cortex and subcortex. Through generating a unit-less value per brain region that describes a pharmacodynamic-informed measure of importance, or strength, of that brain region in the drug’s action relative to all other brain regions, this method also enables their contextualization through neuroanatomical features, such as functional imaging-derived network partitions (15), meta-analytical topic-related activation patterns (16) as well as cytoarchitectural (17) properties. We finally put the distribution profiles in context with imaging-derived morphological changes associated with major depressive disorder to investigate whether antidepressant distribution profiles coincide with disease-associated atrophy patterns (18).

## Results

### Molecular affinity-defined groups of antidepressants and their efficacy

The investigated antidepressants are listed in **Table 1**. From the original study by Cipriani and colleagues, we excluded Levomilnacipran, Vortioxetine and Vilazodone, since KiDB contained no inhibition constants from human sources for these substances.

**Table 1.** List of antidepressant substances. Mean efficacy is derived from a meta-analysis by Cipriani et al. (7). TCA: Tricyclic Antidepressant; SNRI: Serotonin and Noradrenaline Reuptake Inhibitor; SSRI: Selective Serotonin Reuptake Inhibitor; NDRI: Noradrenaline and Dopamine Reuptake Inhibitor; NARI: Noradrenaline Reuptake Inhibitor

| Antidepressant | Abbreviation | Mean efficacy | Substance group |
| --- | --- | --- | --- |
| Amitriptyline | AMI | 2.13 | TCA |
| Mirtazapine | MIR | 1.89 | Atypical |
| Duloxetine | DUL | 1.85 | SNRI |
| Venlafaxine | VEN | 1.78 | SNRI |
| Paroxetine | PAR | 1.75 | SSRI |
| Milnacipran | MIL | 1.74 | SNRI |
| Fluvoxamine | FLUV | 1.69 | SSRI |
| Escitalopram | ESC | 1.68 | SSRI |
| Nefazodone | NEF | 1.67 | Atypical |
| Sertraline | SER | 1.67 | SSRI |
| Agomelatine | AGO | 1.65 | Atypical |
| Bupropion | BUP | 1.58 | NDRI / Atypical |
| Fluoxetine | FLUO | 1.52 | SSRI |
| Citalopram | CIT | 1.52 | SSRI |
| Trazodone | TRA | 1.51 | Atypical |
| Clompiramine | CLO | 1.49 | TCA |
| Desvenlafaxine | DVEN | 1.49 | SNRI |
| Reboxetine | REB | 1.37 | NARI |

For the remaining substances, we employed hierarchical agglomerative clustering and k-means clustering analyses on log-scaled KiDB-derived molecular affinity profiles to develop a data-driven systematization of antidepressants (**Figure 1A, B**). Unlike hierarchical agglomerative clustering, k-means clustering requires a previously selected number of centroids that govern the resulting number of clusters. We sampled the parameter space of centroids and used silhouette coefficients to assess the optimal cluster number, which we determined to be three (**Supplementary Figure 1**). Sorting of substances into different clusters matched between hierarchical agglomerative clustering and k-means clustering. Of the three clusters, the first cluster contained Amitriptyline and Clomipramine, the second cluster contained Mirtazapine, Nefazodone and Trazodone, and the third cluster was composed of the remaining substances. To ensure that the formation of these clusters were not simply due to the tuning of the clustering algorithms, we replicated the analysis using different linkage algorithms and data transformations in hierarchical agglomerative clustering, were they presented as stable (**Supplementary Figures 2 and 3**).

**Figure 1.**
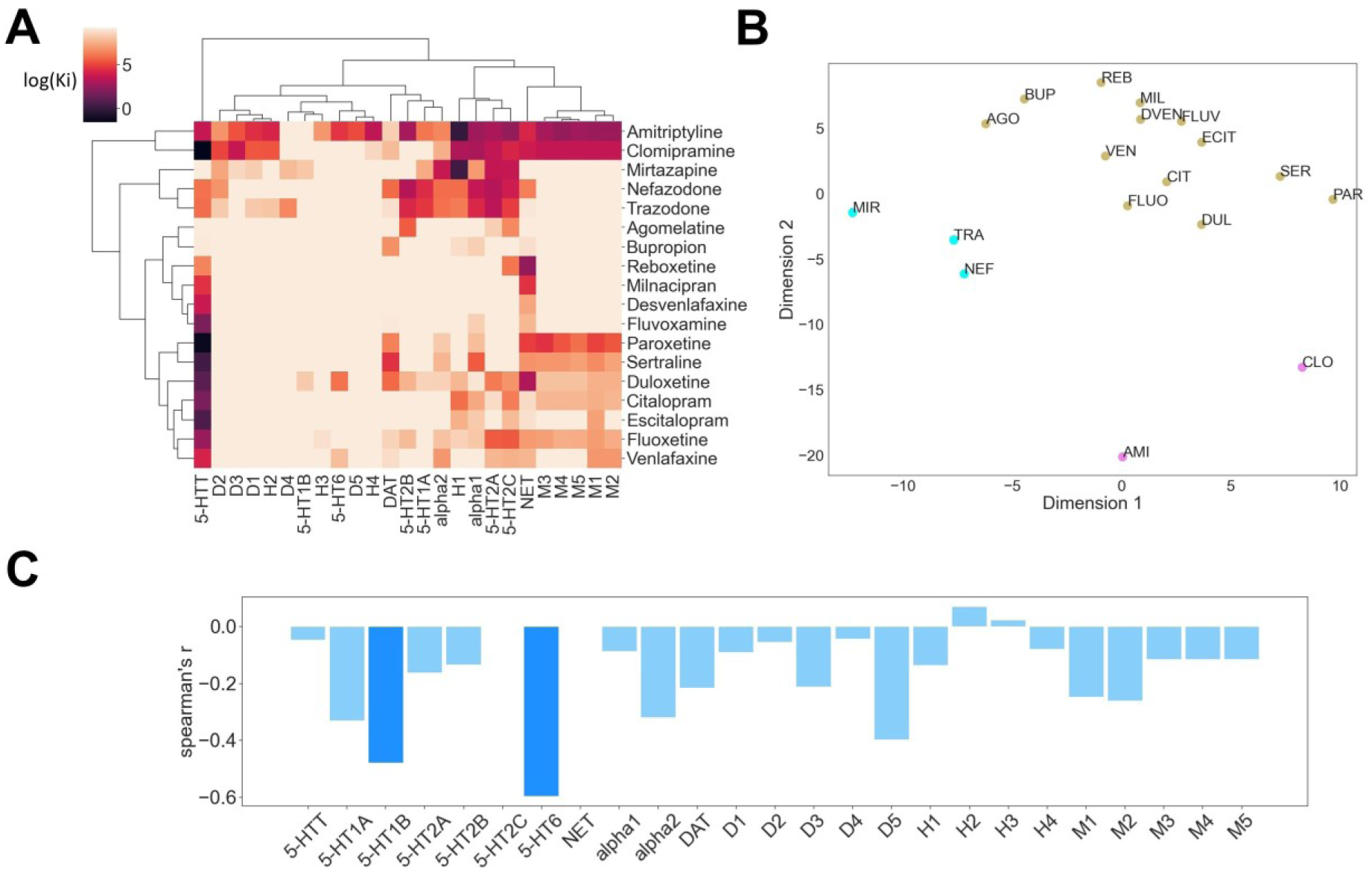
Molecular affinity profiles of antidepressants. **A)** Hierarchical agglomerative clustering of log-scaled KiDB-derived inhibition constants across all substances. **B)** MDS-derived two-dimensional scatter plot of k-means clustering results of the same inhibition constants. Scatter colours denote cluster memberships. **C)** Spearman rank correlation of receptor and transporter affinities and antidepressant efficacies across all substances. Saturated blue colour represents nominally statistically significant correlations at *p < 0.05*, of which none survived controlling for multiple comparisons.

Based on our clustering results, we proceeded to investigate whether the substance clusters differed in their antidepressant efficacy and found no significant difference in mean antidepressant efficacy between the clusters (**Supplementary Figure 4A**). As affinity-based clusters showed to be non-informative regarding antidepressant efficacy, we investigated whether there were significant associations between molecular affinities and antidepressant efficacy through correlation analysis. Across all substances, we found nominally significant negative correlations with antidepressant efficacy which did not survive false discovery control at *⍺* = 0.05 for 5-HT1B (corrected p-value = 0.38) and 5-HT6 (corrected p-value = 0.12) affinities (**Figure 1C**). As lower inhibition constants signify greater affinity, the negative relationship indicates that higher affinity to these receptors and greater antidepressant efficacy coincide. Notably, while most efficacious antidepressants show their strongest affinity to 5-HTT, no significant correlation between 5-HTT and antidepressant efficacy was found, neither across all substances, nor in the biggest cluster that encompasses serotonin reuptake inhibitors (**Supplementary Figure 4B**).

### Cortical antidepressant distribution profiles

In the next step, we investigated the topography of antidepressant distribution in the human cerebral cortex, which necessitates both the underlying cerebral chemoarchitecture, as well as the antidepressants’ molecular affinity profiles. We assessed spatial maps of chemoarchitecture through an open-access collection of PET-derived neurotransmitter transporter and receptor density maps (14), which are reported on in **Table 2**. Subsequently, for each drug, we calculated a parcel-wise weighted average of neurotransmitter transporter and receptor density values, with the drug’s respective molecular affinity profile governing the weighting (**Methods, Equation 1**). The resulting value is a measure for the drug’s regional receptor/transporter binding or occupancy, and thus a proxy measure for localized drug action strengths that allows for inter-regional comparisons. Cortical projections of the most efficacious antidepressants per previously defined cluster are displayed in **Figure 2A**, cortical projections of the other substances ordered by clusters are shown in **Supplementary Figure 5A.** To further contextualise the anatomical patterns, we quantified the distributions of drug action values across resting-state fMRI derived functional networks (15) and a framework of cytoarchitectural classes that links cortices’ histological compositions to the supposed complexity of cognitive functions they realise (17) **(Figure 2B)**.

**Figure 2.**
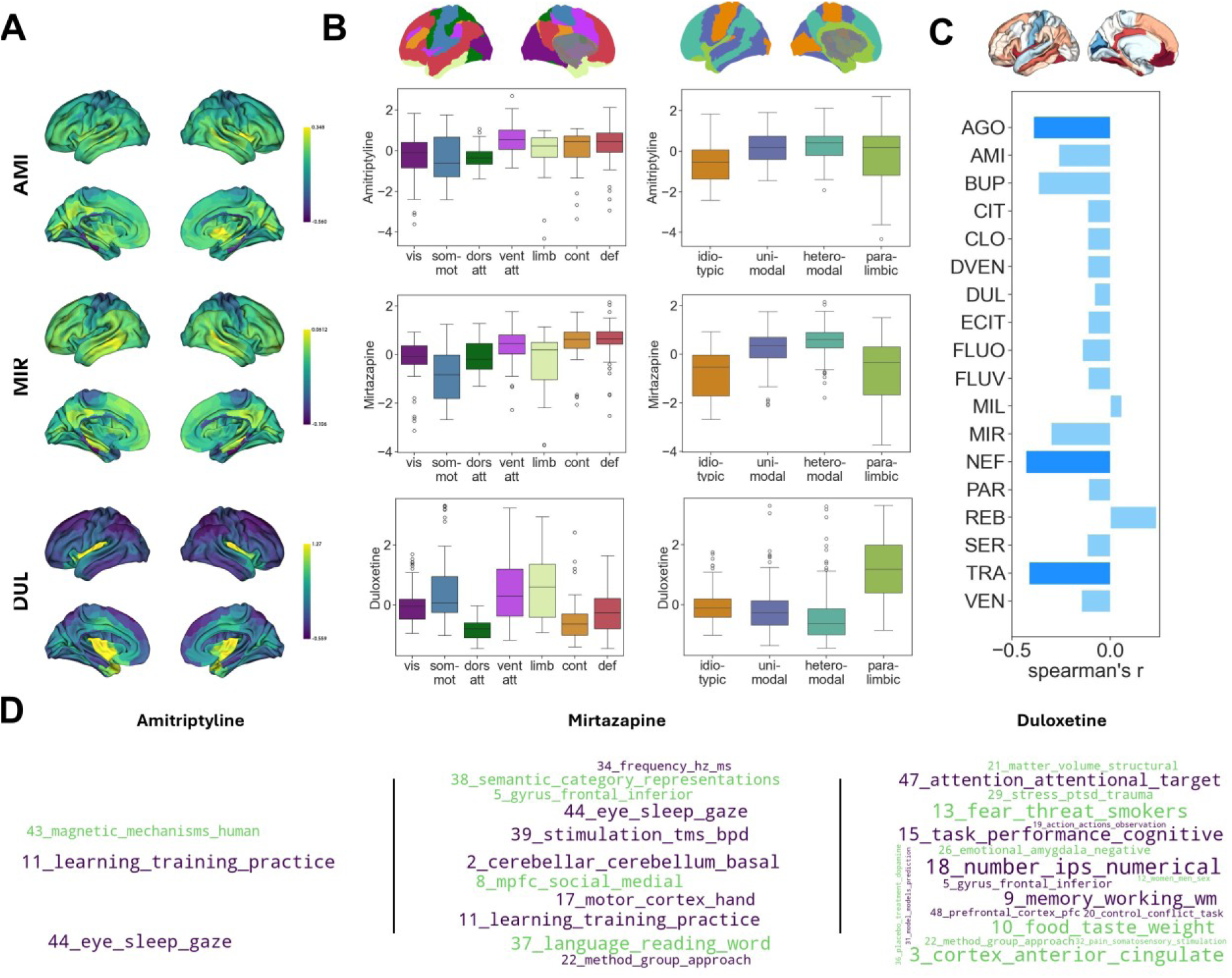
Cortical antidepressant distribution profiles. **A)** Cortical projections of Amitriptyline, Mirtazapine and Duloxetine drug action values, which are the most efficacious antidepressants in their respective clusters. **B)** Anatomical contextualization of Amitriptyline (top), Mirtazapine (middle) and Duloxetine (bottom) cortical distribution profiles. Distribution of drug action values across functional networks (left) and cortical types (right). **C)** Spearman rank correlation of cortical antidepressant distribution profiles with disease-associated cortical thickness alterations in major depressive disorder. Top: Cortical projection of disease-associated cortical thickness alterations. Bottom: Bar plot of correlation strengths. Saturated blue colours represent nominal statistically significant correlations at *p < 0.05*. **D)** Term-based meta-analytical decoding of cortical antidepressant distribution profiles. Terms with nominally statistically significant correlations (*p < 0.05)* are displayed. Green indicates positive correlations, purple indicates negative correlations, and font size indicates correlation strength.

**Figure 3.**
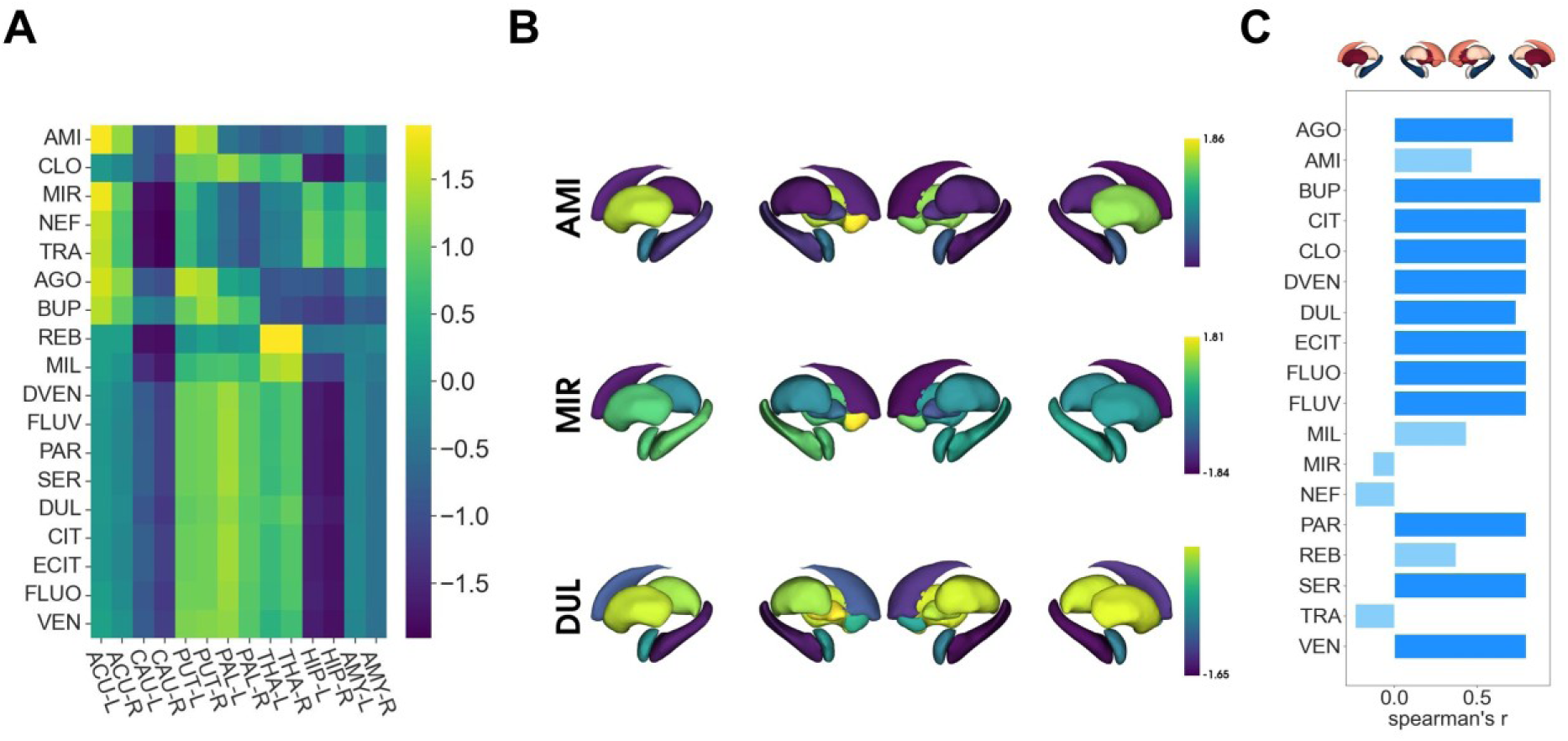
Subcortical antidepressant distribution profiles. **A)** Heatmap display of drug action values in subcortical structures across substances (ACU – Accumbens nucleus; CAU – Caudate nucleus; PUT – Putamen; PAL – Pallidal globe; THA – Thalamus; HIP – Hippocampus; AMY – Amygdala). **B)** Subcortical projection of drug action values of Amitriptyline (top), Mirtazapine (middle) and Duloxetine (bottom). **C)** Spearman rank correlation of subcortical antidepressant distribution profiles with disease-associated subcortical volume alterations in major depressive disorder. Top: Subcortical projection of depression-associated volume aberrations. Bottom: Bar plot of correlation strengths. Saturated blue colours represent nominal statistically significant correlations at *p < 0.05*.

**Table 2.** List of PET-derived neurotransmitter transporter and receptor molecules. Regular font designates receptors, italic font designates transporters. The table is adapted from (52). For further information on the neurochemical maps, including sample sizes, age and sex distributions, and details on the respective PET scans, see (14).

| Transporter / Receptor | Abbreviation | Tracer | References |
| --- | --- | --- | --- |
| Dopamine receptor D1 | D1 | [11C]SCH23390 | (61) |
| Dopamine receptor D2 | D2 | [11C]FLB-457 | (62,63) |
| *Dopamine transporter* | *DAT* | [123I]-FP-CIT | (64) |
| *Norepinephrine transporter* | *NET* | [11C]MRB | (65) |
| 5-HT1A receptor | 5-HT1A | [11C]WAY-100635 | (55,66) |
| 5-HT1B receptor | 5-HT1B | [11C]P943 | (66,67) |
| 5-HT2A receptor | 5-HT2A | [11C]Cimbi-36 | (55) |
| 5-HT6 receptor | 5-HT6 | [11C]GSK215083 | (55,68) |
| *5-Hydroxytryptamine transporter* | *5-HTT* | [11C]DASB | (55) |
| Muscarinic acetylcholine receptor M1 | M1 | [11C]LSN317217<br>6 | (69) |
| Histamine H3 receptor | H3 | [11C]GSK189254 | (70) |

Antidepressants displayed nuanced cortical distribution patterns across molecularly defined clusters, although correlational analyses showed a notable across-cluster influence of the degree of 5-HTT affinity with regards to distribution profile similarity **(Supplementary Figure 5B)**. Amitriptyline, representing the first cluster, showed a comparatively homogeneous distribution profile across the cerebral cortex. Higher drug action values were apparent in the anterior insular cortex and the parietal lobe. Decoding analyses showed lower drug action values in visual and somato-motor networks and idiotypic cortices, and higher values in the ventral attention and default mode networks and uni- and heteromodal cortices. Decoding analyses of Mirtazapine, representing the second cluster, revealed similar trends, but in a more pronounced fashion. Here, the difference in drug action values between somato-motor and default mode networks was higher, and drug action strengths increased across a gradient from idiotypic to heteromodal cortices. Duloxetine, representing the third cluster, showed high values in the entorhinal and insular cortices and the temporal lobe, where the temporal pole presented particularly prominent (**Figure 2A, Supplementary Figure 5A**). Likewise, in the decoding analyses, its drug action values were high in the limbic functional network and in paralimbic cortices.

Furthermore, we investigated whether the distribution profiles of antidepressants coincide with depression-specific alterations in brain morphology which may reflect disease-associated neuroplastic changes (19,20). To this end, we performed correlation analyses between antidepressant distribution profiles and cortical thickness alterations in patients with major depressive disorder from a large multi-site sample (18). Both measures were surface-projected, and spin-permutation tests were used to assess statistical significance (see **Methods**). Notably, the cortical distribution profiles of Agomelatine, Trazodone and Nefazodone showed nominally significant negative correlations to cortical thickness changes associated with major depressive disorder, of which none survived false discovery control at *⍺* = 0.05 (corrected p-values = 0.117). The negative correlations indicate a positive spatial relationship between antidepressant action strength and disease-associated cortical thinning (**Figure 2C**).

As the anatomical and disease-associated decoding analyses hinted at differential relationships of antidepressant distribution profiles to underlying structure-function relationships in the brain, we opted to further dissect their relationship to functional anatomy. As such, we employed topic-associated meta-analytical activation maps generated from a large repository of fMRI studies (21) to assess their relationships to localized functional activations associated with relevant concepts in cognitive neuroscience. Amitriptyline showed significant positive correlations with a technical topic without clear functional implication (topic 43). Mirtazapine showed significant positive associations with topics related to language and representation-related higher-order cognitive functions (topics 37,38) and social reasoning and theory of mind (topic 8). Duloxetine correlated positively and significantly with topics of limbic anatomy (topic 3) and function, as in homeostatic signaling (topic 10), fear and threat conditioning (topics 13, 29) and emotion processing (topic 26) (**Figure 2D**).

### Subcortical antidepressant distribution profiles

In a final step, we generated antidepressant distribution profiles in subcortical structures (**Figure 3A, B**). Nucleus accumbens shows high drug action values for Amitriptyline, Agomelatine, Bupropion, and atypical antidepressants, and low values for SSRI, SNRI and Clomipramine. All substances show low drug action values in the Caudate nucleus, and high values in Putamen. Pallidum and Thalamus show high drug action values for SSRI, SNRI, Bupropion and Clomipramine, while atypical antidepressants and Amitriptyline show low values these nuclei. High drug action values found in Hippocampus and Amygdala stem from Mirtazapine, Nefazodone and Trazodone, while the other substances show low values in the two structures.

Similarly to our cortical analysis, we investigated the relationship between drug distribution profiles and depression-specific alterations in the volume of subcortical nuclei. Except for Amitriptyline, Milnacipran, Mirtazapine, Nefazodone, Reboxetine and Trazodone, the distribution profiles showed nominally and multiple comparisons-corrected significant positive correlations with depression-associated alterations in subcortical volume. Significance was established using permutation testing against spatial autocorrelation-preserving surrogate maps generated using variogram matching (see **Methods**). The correlation sign implies a negative spatial relationship between disease-associated subcortical atrophy and high drug action strength **(Figure 3C)**.

### Stability analyses

To address potential effects of parcellation granularity, we replicated the cortex-based analyses using the 100 parcel Schaefer parcellation **(Supplementary Figure 6)**. Except for the correlation between drug distribution profiles and disease-associated cortical thickness alterations losing nominal statistical significance while approximately retaining correlation strengths, the results are stable overall.

## Discussion

In this study, we present a data-driven classification of antidepressants intro three distinct clusters based on their multidimensional molecular affinity profiles. Cluster membership and individual molecular affinities were not significantly associated with antidepressant efficacy. Next, we establish cortical and subcortical distribution maps of antidepressants based on their affinities and the brain’s chemoarchitecture. We then investigate the relationship between the cortical maps with functional and cytoarchitectural neuroanatomical measures. Our results reveal distinct cluster-associated patterns. Furthermore, we demonstrate that two groups of antidepressants exhibit opposing relationships with disease-associated morphological alterations. Atypical antidepressants correlate with major depressive disorder-associated cortical thickness alterations, while antidepressants with high 5-HTT affinity correlate with subcortical volume alterations. In summary, our results demonstrate the affinity-based systematization and context-imbued cortical and subcortical projection of clinically efficacious antidepressants.

Our affinity profile-derived taxonomy recapitulates both the group of tricyclic antidepressants (Amitriptyline, Clomipramine) in the first cluster, as well as the group of atypical antidepressants (Mirtazapine, Nefazodone, Trazodone) in the second cluster. Agomelatine, which is exempt from the atypical cluster, differs from the other substances as it is a potent agonist at melatonin receptors, for which neither inhibition constants nor distribution data were available. The remaining third cluster, however, intermixes traditionally separate groups of different reuptake inhibitors, and no stable further separation was sensibly attainable. Our data-driven division between antidepressant groups is thus considerably coarser than the traditional taxonomy. This finding might be important in the context of studies that investigate the switching of antidepressant medication in case of a clinical non-response. Current evidence regarding the switching of medication within or between the traditional antidepressant groups is inconclusive, which is also reflected in variations in treatment recommendations (22,23). Our clustering and correlation results hint that the neurochemical profiles targeted by SSRI and SNRI respectively might not be as strikingly different as could be expected, which is strengthened by our neuroanatomical evidence showing a strong correlation of cortical distribution profiles between SSRI and SNRI. Future studies investigating the best course of action after pharmacological non-response might take such revealed similarities into consideration.

The affinity profile-derived clusters were non-informative regarding antidepressant efficacy, and they revealed no cluster-specific receptor or transporter affinities that significantly relate to efficacy. Across all substances, our study found only nominally significant associations between 5-HT1B and 5-HT6 affinities and antidepressant efficacy. Mirtazapine and Duloxetine show 5-HT1B affinity, while Amitriptyline, Duloxetine and Venlafaxine show 5-HT6 affinity, and these substances are the four most effective antidepressants in the meta-analysis by Cipriani and colleagues (7). Both of these metabotropic serotonin receptors have been studied in the context of antidepressant treatment. A 5-HT6 polymorphism was associated with increased response to antidepressant treatment (24), and both selective agonists and antagonists could show antidepressant-like responses in animal models (25), including a potentiation of sub-therapeutic doses of antidepressants through 5-HT6 antagonists (26). Biochemically, antagonizing 5-HT6 was associated with an increased release of monoamines in rats (27). 5-HT1B is an autoreceptor that inhibits serotonin release upon activation (28), and its downregulation was associated with antidepressant treatment both in animal models and in humans (29). Analogously to 5-HT6, combining antidepressants with 5-HT1B antagonists showed a potentiated antidepressant effect in animal models (30). While our results similarly imply 5-HT1B and 5-HT6 as potentially relevant targets in antidepressant actions, the strength of our evidence may be limited - the neurobiological effects the receptors mediate under current antidepressant treatments may be questionable, as the inhibition constants for 5-HT1B and 5-HT6 exceed the expected drug concentrations, except in the case of Amitriptyline (31).

The cortical and subcortical distribution maps of antidepressants constitute a novel approach to investigating pharmacodynamic properties in the neuroimaging domain. Studying psychopharmacologic drugs in anatomical imaging spaces may constitute a first step at bridging findings in basic neurosciences as well as disease-associated imaging alterations with pharmacological interventions. Regarding the cortex, the dominance of antidepressants with considerable 5-HTT affinity in (para)limbic areas allows for intriguing hypotheses about the drugs’ mechanism of action through the functional properties of these regions, which show prominent connections to subcortical and prefrontal areas (32) and are thought to be more prone to neuroplastic changes due to their histological composition (33). Generally, the complex cognitive functions realized in (para)limbic areas (34,35) hold relevance in psychiatric illnesses. Especially the high action values in the temporal pole and entorhinal and insular cortices hold interesting implications. The temporal pole is hypothesized to link emotional responses to highly processed sensory input (36), and temporal pole atrophy was found in multiple psychiatric diseases from the affective spectrum and in OCD and anxiety patient groups (37). The insula, a site of cross-disorder grey matter atrophy in psychiatric disorders (38), is involved in emotion recognition (39). Indeed, our functional decoding analysis showed relationships to terms associated with emotion processing and recognition. Given this underlying functionality, medication-induced alterations in temporopolar and insular cortices could plausibly impact the processing of emotionally salient stimuli, which would be in accordance with neuropsychological theories of antidepressant action (6). The entorhinal cortex, on the other hand, has been implicated as a key driver of hippocampal neurogenesis (40). Strong antidepressant action in this region could thus mediate the neurogenetic effects at the heart of the neuroplasticity theory of antidepressant action (5). Here, the subcortical drug distribution profiles, which show little activity of 5-HTT-affine substances in the hippocampus, further support claims towards their indirect rather than direct effects on hippocampal neurogenesis (41). We therefore claim and demonstrate that non-exclusive theories of antidepressant action from different neuroscientific disciplines can be sensibly linked through integrating pharmacodynamic properties into the neuroimaging space. Furthermore, especially antidepressants with a dominant 5-HTT affinity showing strong action values in regions which are central to emotion processing may explain why they are not only effective in treating depression, but also other psychiatric disorders like obsessive-compulsive disorder and anxiety disorders, to both of which strong unpleasant emotions are central.

Furthermore, we detect correlations between Agomelatine, Trazodone and Nefazodone distribution profiles and cortical thinning patterns found in major depressive disorder. These antidepressants share that their affinity profiles are not 5-HTT dominated, and that they are traditionally classified as atypical. Cortical thinning - which can constitute a proxy measure for neuronal and synaptic density (42) - in adult major depressive disorder is pronounced in areas that interact closely with the limbic system (18), encompassing cortical areas located primarily in the limbic and default mode networks. These abnormalities show closer correspondence to the comparatively less localized distribution profiles of the non-5-HTT-dominant antidepressants, opposed to the more localized distribution profiles of SSRI, SNRI and Clomipramine. Interestingly, in the subcortex, SSRI, SNRI and Clomipramine show positive correlations to disease-associated changes in subcortical volume, of which the coinciding low drug action values and volume reduction in the hippocampus are likely drivers (43).

Summarised across all analyses, our results hint at the existence of two distinct pharmacodynamically driven mechanisms of action between 5-HTT-dominant and atypical antidepressants. In neuroanatomical contextualization analyses, atypical antidepressants showed stronger values in heteromodal cortices and default mode network regions, which plausibly realise language and representation-related higher-order cognitive functions and social reasoning and theory of mind detected in the meta-analytical decoding analyses of atypical antidepressant’s drug distribution profiles. 5-HTT-dominant antidepressants, on the other hand, act strongly in cortices of paralimbic cytoarchitectural differentiation and limbic functionality, which again plausibly realise the functions of homeostatic signaling, fear and threat conditioning and emotional processing detected in meta-analytical decoding. Furthermore, the opposing relationships of atypical substances and drugs with high 5-HTT affinity to disease-associated morphology changes in the cortex and subcortex may hint at two different morphology-associated mechanisms of action. From a network perspective on depression (44), the atypical group would induce changes in multiple nodes of a morphological depressive network, where they functionally primarily alter highest-order cognitive functions like theory of mind, social reasoning and top-down emotion regulation. Substances with high 5-HTT affinity, on the other hand, would act primarily at the cortical hubs of limbic functionality, where they functionally impact the emotional processing of complex stimuli and homeostatic signals. These findings might inform future studies that investigate whether structural morphological alterations on the individual patient level can be leveraged to inform and further personalize the choice of antidepressants (45), and likewise investigations that seek to connect symptom profiles to choice of antidepressants.

There are several important limitations to consider when interpreting this study, pertaining first to the KiDB-derived molecular affinities. First, the inhibitory constants from KiDB varied across studies, which we addressed through averaging the values. Even though we restricted ourselves to human samples, some uncertainty remains in how far the Ki values used in this study represent pharmacodynamic interactions in the human brain. Regarding the neuroanatomical findings, the antidepressant distribution maps were generated using chemoarchitectural data from healthy participants, which might differ in individuals suffering from major depressive disorder, as especially medication-induced alterations in receptor and transporter expression cannot be accounted for (46). It is furthermore important to consider that, in calculating the distribution profiles, we can only consider relative, but not absolute protein densities due to the PET resource largely reporting Binding Potential (BPnd) and Standardised Uptake Value Ration (SUVR) measures, which are not directly comparable to one another. Therefore, the contribution of sparsely expressed proteins to the distribution profiles might be over-inflated, while the contribution of densely expressed proteins might be under-weighted. As estimating absolute regional protein density measures with PET imaging is challenging (47), this might be considered an inherent limitation of the method that is exacerbated when aiming to cover a multitude of different protein densities. Once available, however, future studies of drug distribution profiles should leverage absolute protein density measures.

Furthermore, the overlap between KiDB-reported affinities and available PET-derived neurotransmitter transporter and density maps is only partial. The particular absence of alpha1 and alpha2 receptor distribution maps has to be pointed out, as no suitable radiotracers for imaging these receptors in humans are available yet (48). This is important when looking at the distribution profiles of the relatively alpha-affine atypical substances Mirtazapine, Nefazodone and Trazodone.

Additionally, the antidepressant efficacy drawn from the extensive meta-analysis by Cipriani et al. pertains to a > 50% symptom reduction in major depressive disorder. The efficacy data has to be viewed critically due to multiple considerations regarding the interpretation of the original study (for a detailed comment, see (49)), and the overall difference in efficacy is low, which the authors in the original meta-analysis also conclude. Notwithstanding these considerations, we still view it as one of the best current sources for overall antidepressant efficacy and hence employ its findings while keeping the caveats in mind. Finally, we point out that multiple statistical significances in our correlational analyses did not survive the correction for a false discovery rate at *⍺* = 0.05 using the Benjamini-Hochberg method. While this naturally implies that the significance of the correlational analyses might be spurious, there are ongoing debates questioning the merit of controlling for multiple comparisons in exploratory studies of this nature (50). For full transparency, we therefore chose to report on both nominal as well as adjusted significance and caution the reader to keep this distinction in mind.

In conclusion, our results suggest that, when looking at putative relationships between pharmacodynamic properties and mechanisms of action, the multiplicity of antidepressant substances can be reduced to an even sparser set than traditionally considered. This is reflected both in our clustering-based as well as the neuroanatomically-based results and holds important implications for future drug development and personalized psychiatry efforts. The latter are especially crucial to further understand the heterogeneity of major depressive disorders across patient collectives with vastly different characteristics. Dissecting the intricacies of the disorder will be crucial to improve its treatment and lessen both its personal as well as its societal burden. Finally, our approach to studying psychopharmacological substances in imaging spaces lets us discern sensible connections between functional neuroanatomical features and working theories of antidepressant action. This approach can be expanded towards other drugs and disorders (51), providing a novel basis to link imaging and neuroanatomical findings to psychopharmacological research in future studies.

## Methods

### KiDB

To generate molecular affinity profiles, we sourced inhibition constants from KiDB, an open-access repository that collects published inhibition constants (13). Only Ki values from sources flagged as ‘human’ were used. For each substance-transporter/receptor interaction, the reported Ki values were averaged across studies after excluding studies that reported Ki values inexactly (e.g., specifying a Ki value of > 10000). Ki values were capped at 10000, which was also used to represent missing data. Three substances from the original meta-analysis (Vortioxetine, Vilazodone, and Levomilnacipran) were excluded due to insufficient inhibitory constants being present in the database. The final extract of KiDB is presented in **Supplementary Table 1.**

### Clustering

Two different clustering techniques on the KiDB-derived molecular affinity profiles were performed: Hierarchical agglomerative clustering, using euclidean distance and Ward’s linkage algorithm, and k-means clustering. To select the number of centroids in k-means clustering, we used silhouette coefficients, defined as 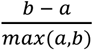, where *a* is the mean intra-cluster distance and *b* is the mean nearest-cluster distance. Clustering analyses were replicated using native and log-transformed Ki values. Log-transformation was performed to ensure that distances between compounds reflect relative differences in binding affinity, since Ki measurements are approximately log-normally distributed.

### PET data preparation and antidepressant distribution profile generation

To assess neurotransmitter transporter and receptor molecule (NTRM) densities in the brain as a prerequisite for generating cerebral antidepressant distribution profiles, we leveraged a collection of open-access PET maps from healthy participants (14). We performed PET data preparation similar to a previous description (52) - briefly, the PET density maps were parcellated to 400 regions based on the Schaefer parcellation (53) to obtain cortical parcels, and subcortical regions were extracted using freesurfer, or aseg-based, subcortical regions of interest (54). Following original author recommendations, we extracted data from surface-projected receptor density maps for 5-HTT, 5-HT1A, 5-HT2A and 5-HT6 (55). Weighted averages were generated when multiple studies per radiotracer were available. The resulting parcel-wise or subcortical regions-wise intensity values were z-score normalized per tracer, where cortical and subcortical compartments were treated separately.

To subsequently calculate the anatomical antidepressant distribution profiles, the following receptors and transporters were used to calculate the cortical antidepressant distribution profiles, as both inhibition constants as well as PET density maps were available for them: 5-HTT, NET, DAT, 5-HT1A, 5-HT1B, 5-HT2A, 5-HT6, D1, D2, M1, H3. For each substance, a unit-less value *a* per region was calculated as

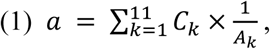

with *C* and *A* being one-dimensional arrays representing the same NTRM’s density (*C*) and Ki value (*A*) at location *k*. For each parcel, the value *a* represents the expected strength of drug action based on that region’s chemoarchitecture and the affinity profile of the drug in question.

### Functional network and cytoarchitectural classes decoding

To contextualize cortical antidepressant distribution profiles, we leveraged a resting-state fMRI-based partition of the cortex into seven functional networks, ‘Yeo networks’ (15) and a theoretical framework of cytoarchitectural classes that describe a sensory-fugal gradient (17).

### ENIGMA data

To probe associations between disease-associated alterations in cerebral morphology and antidepressant distribution profiles, we employed multisite summary statistics of cortical thickness and subcortical volume variation published by the ENIGMA Consortium (56) that are publicly available through the enigma toolbox python package (57). Cortical thickness and subcortical volume alterations were reported in the form of Cohen’s *d*, denoting covariate-adjusted case-versus-control differences in across-site random-effects meta-analyses. The raw data was handled and preprocessed at each site according to standardized ENIGMA quality control protocols (see http://enigma.ini.usc.edu/protocols/imagingprotocols). Summary statistics were derived from adult samples.

### Meta-analytical functional decoding

To assess the relationship between cortical drug distribution profiles and localized brain function, we leveraged meta-analytical, topic-based maps of functional brain activation. Using Nimare, we calculated topic-based activation maps of the Neurosynth v5-50 topic release (https://neurosynth.org/analyses/topics/v5-topics-50/), a set of 50 topics extracted from the abstracts in the full Neurosynth database as of July 2018 using Latent Dirichlet Analysis (16). We projected continuous, non-thresholded activation maps to the surface and performed Spearman rank correlations with surface-projected drug distribution profiles.

### Null models and statistical significance testing

To assess statistical significance when comparing surface-projected data, we applied spin permutation (58) to generate randomly permuted brain maps by random-angle spherical rotation of surface-projected data points, which preserves spatial autocorrelation. Vertex values of the medial wall, and vertex values that were either rotated into the medial wall or onto the surface from the medial wall were discarded for each original and permuted map. For subcortical structures, we employed variogram matching to generate spatial autocorrelation-preserving surrogate maps (59). In each approach, we generated 1000 permuted brain maps using the brainspace python package (60) (https://github.com/MICA-MNI/BrainSpace).

## Acknowledgements and funding

We thank Tobias Kaufmann for his insightful comments on previous versions of the manuscript. B.H. acknowledges funding from the Max Planck School of Cognition’s Clinician Scientist programme. S.L.V. acknowledges funding from the Max Planck Society (Lise Meitner excellence programme), the Jacobs Foundation, the Hector Fellow Academy (Research Career Development Award) and the Helmholtz International BigBrain Analytics and Learning Laboratory (HIBALL).

## Code and data availability

All data used in this manuscript is publicly available. KiDB can be accessed at https://pdsp.unc.edu/databases/kidb.php. PET-derived neurotransmitter transporter and receptor density maps can be accessed through https://netneurolab.github.io/neuromaps/. ENIGMA data can be accessed through https://github.com/MICA-MNI/ENIGMA. Meta-analytical decoding data can be accessed at https://neurosynth.org/analyses/topics/v5-topics-50/. The code to reproduce the results in main and supplementary figures is available at https://github.com/bhaenisch/nchem_antidepressants.

## Author contributions

B.H.: Conceptualization, Data curation, Formal Analysis, Investigation, Methodology, Software, Validation, Visualization, Writing – original draft, Writing – review & editing; S.L.V.: Conceptualization, Resources, Supervision, Writing – original draft, Writing – review & editing

## Ethics statement

This study was approved by the ethics committee of the medical faculty, University of Tübingen (Project Number 492/2024BO2). It re-analyses previously published, openly accessible data. No new data was collected. No data used in this study can be attributed to a specific person.

## Supplementary figures

**Supplementary Figure 1.**
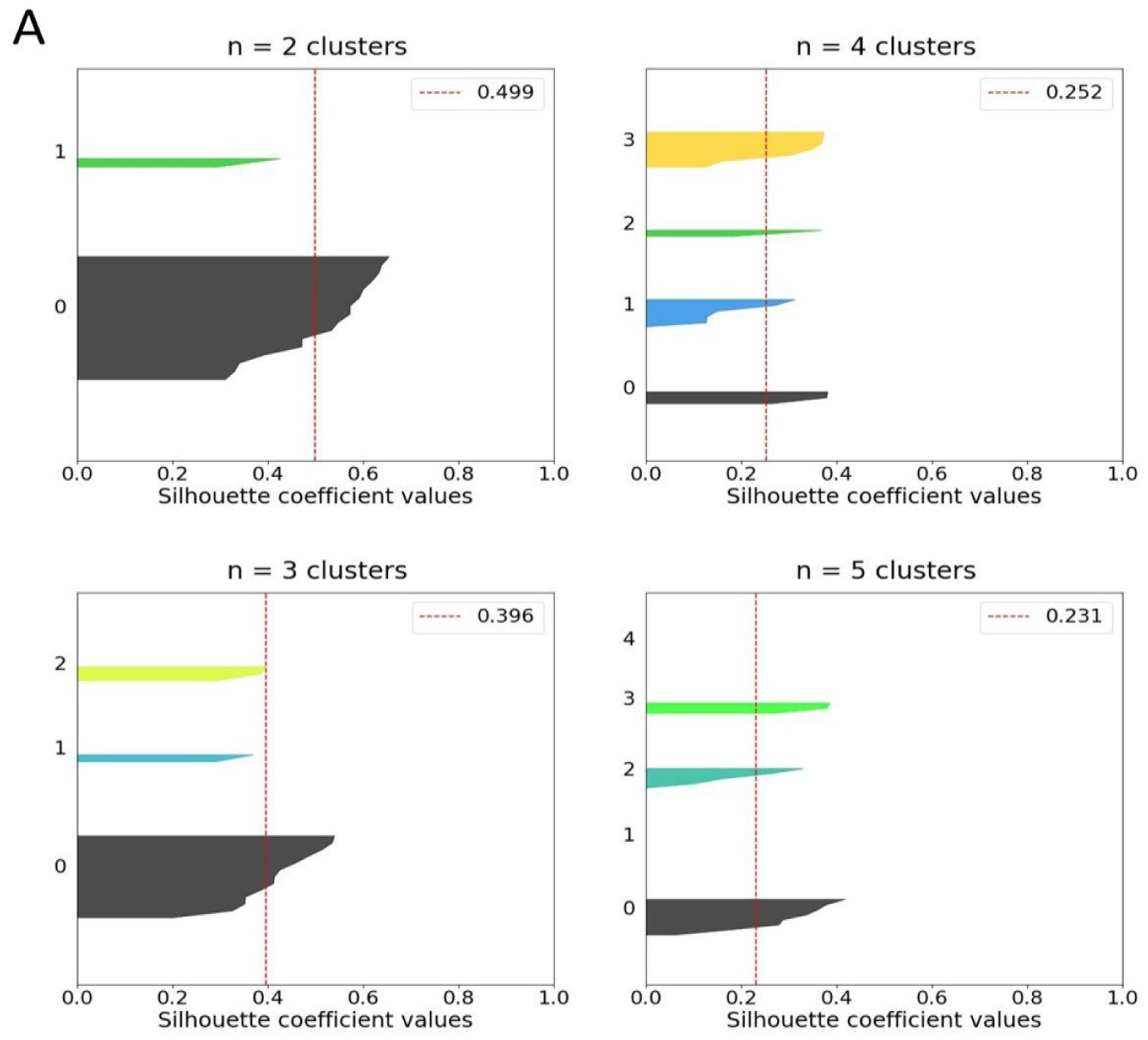
Silhouette coefficients. **A)** Silhouette coefficients of k-means clustering solutions across different pre-specified clustering centroids. Average silhouette coefficient values are displayed as a dashed red line.

**Supplementary Figure 2.**
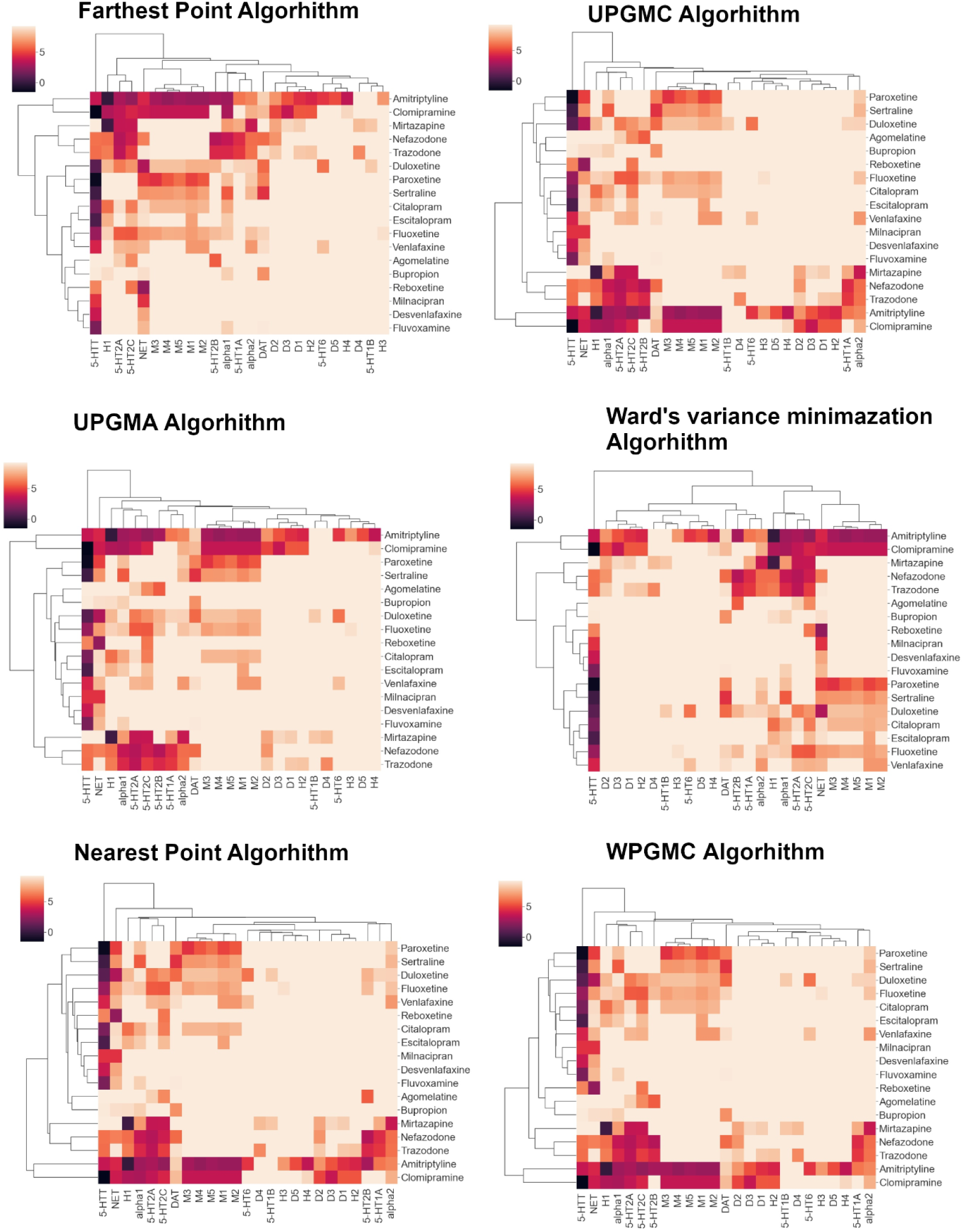
Clustering validation - log transform. Replication of agglomerative hierarchical clustering on log-transformed Ki values using different linkage algorithms.

**Supplementary Figure 3.**
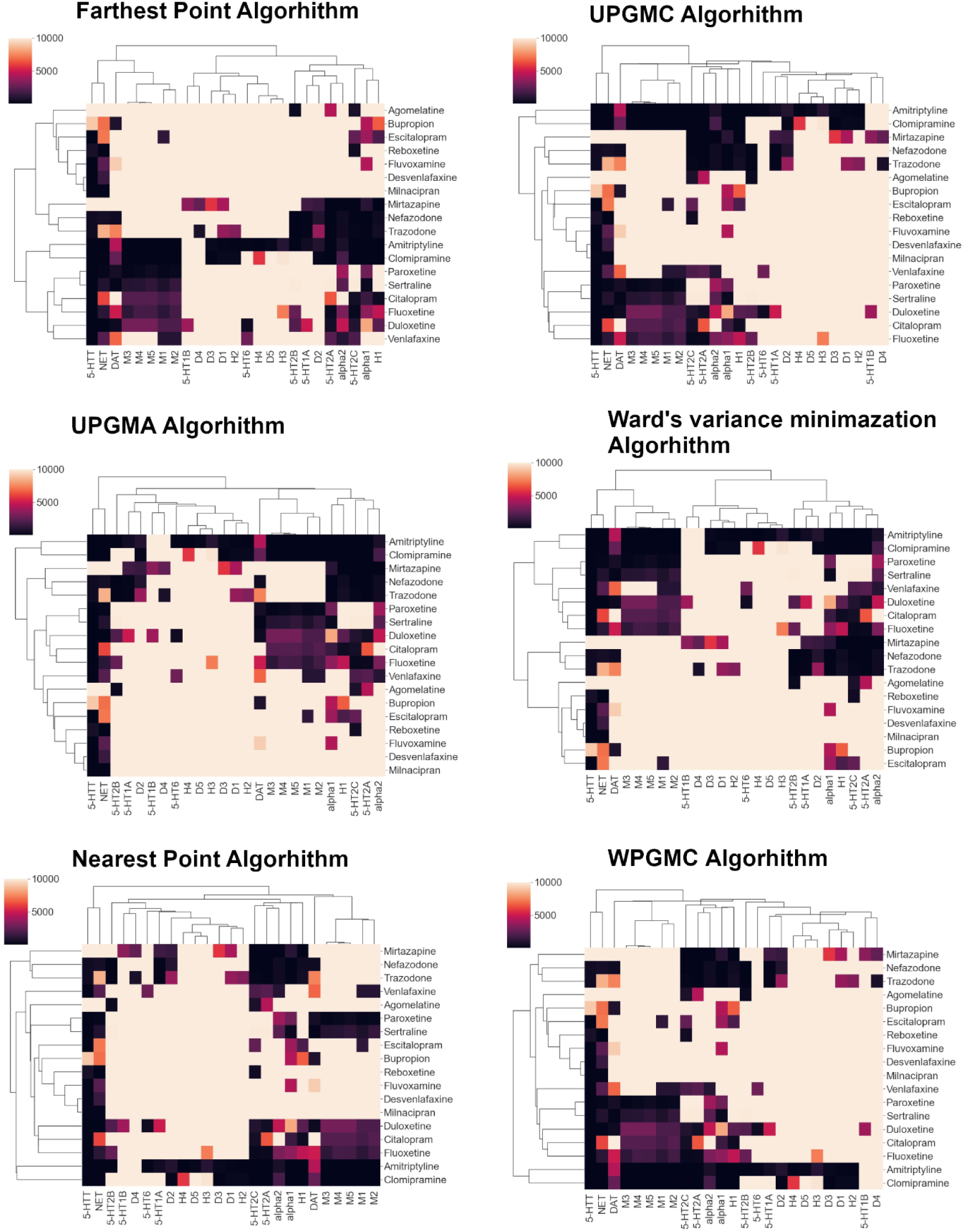
Clustering validation - no transform. Replication of agglomerative hierarchical clustering on non-transformed Ki values using different linkage algorithms.

**Supplementary Figure 4.**
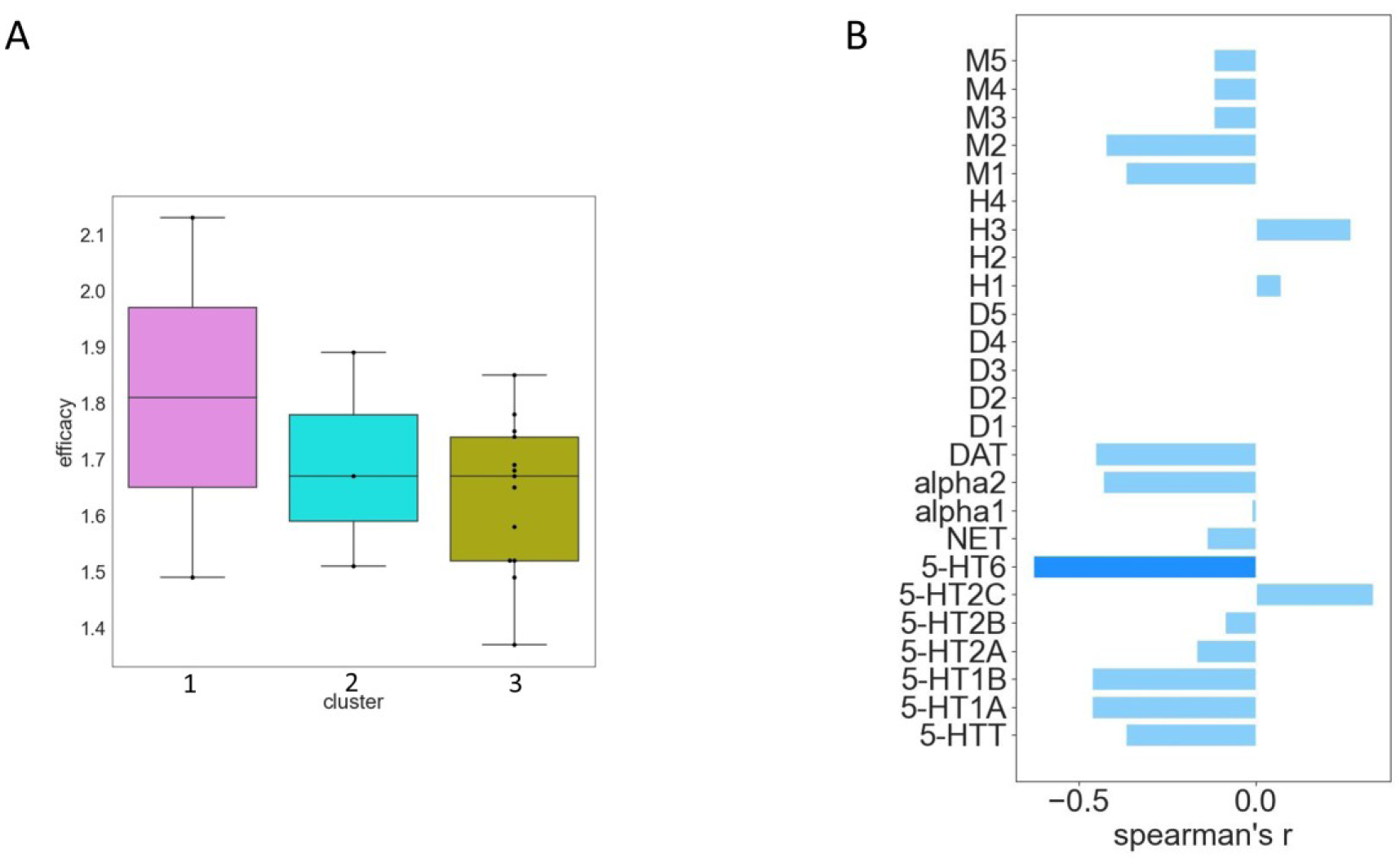
Clustering-based antidepressant efficacies and affinity-efficacy associations. **A)** No significant difference in antidepressant efficacies can be discerned between clusters. **B)** Spearman rank correlations between antidepressant efficacies and affinity profiles for cluster 3. Although this cluster is dominated by 5-HTT affine antidepressants, the degree of 5-HTT affinity is not significantly correlated with antidepressant efficacy.

**Supplementary Figure 5.**
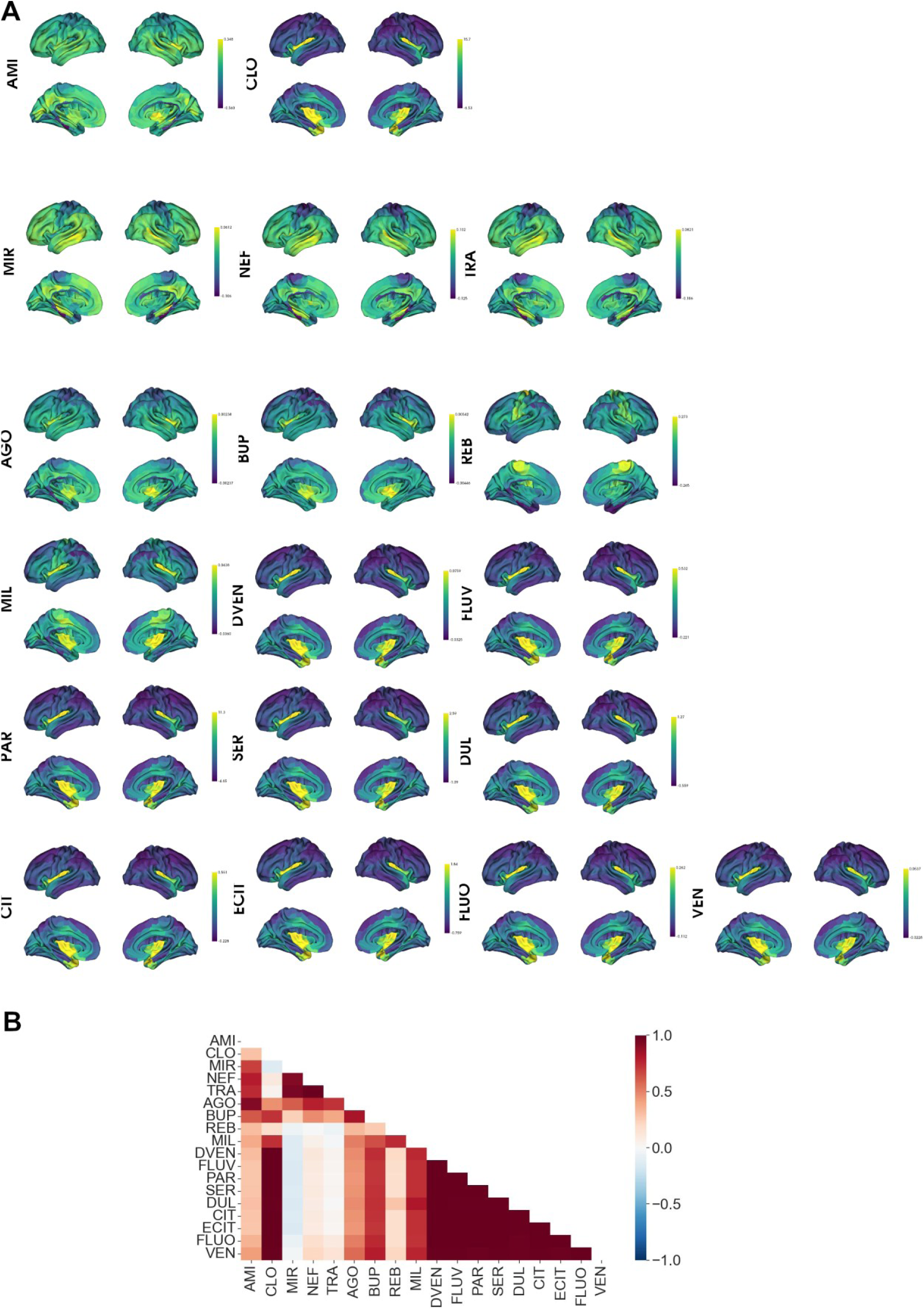
Cortical antidepressant distribution profiles. **A)** Cluster-based cortical projections of antidepressant distribution profiles **B)** Spearman rank correlations between antidepressants’ cortical distribution profiles. Notice that Clomipramine, which has a high 5-HTT affinity, correlates strongly with SSRI and SNRI.

**Supplementary Figure 6.**
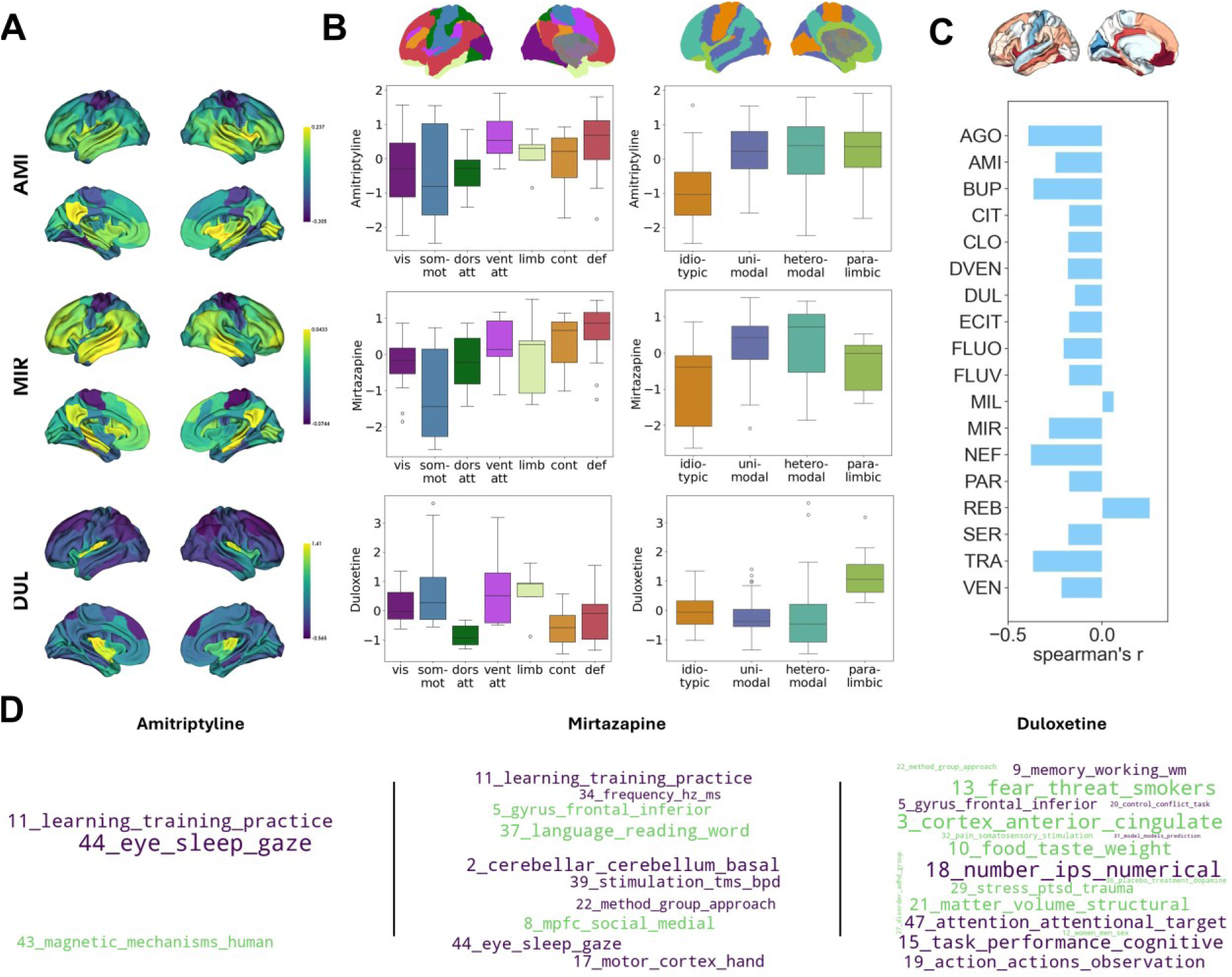
Parcellation stability replication. Replication of Figure 2 using a granularity of 100 Schaefer parcels.

